# In Utero Exposure to Anti-Caspr2 Antibody Disrupts Parvalbumin Interneuron Function in the Hippocampus

**DOI:** 10.1101/2025.04.25.650362

**Authors:** Ciara Bagnall-Moreau, Joshua Strohl, Benjamin Spielman, Christian Cruz, Patricio Huerta, Lior Brimberg

**Author notes:** **Corresponding Author**: Lior Brimberg, PhD, Address: The Feinstein Institutes for Medical Research, 350 Community Dr. Manhasset NY 11030.

## Abstract

Autism Spectrum Disorder (ASD) is a complex neurodevelopmental condition characterized by deficits in communication and social interaction and may stem from an imbalance between excitatory and inhibitory (E/I) signaling in neural circuits. Parvalbumin-expressing (PV+) interneurons are crucial for maintaining E/I balance and regulating network oscillations. Alterations in the number of PV+ interneurons or reductions in PV expression have been observed in both the postmortem brains of individuals with ASD and in animal models, including those induced by in utero exposure to maternal brain-reactive antibodies. In this study, we investigate the impact of in utero exposure to maternal anti-Caspr2 IgG on PV+ interneuron development and function in the hippocampus. Our results demonstrate a selective reduction in PV+ interneurons and perisomatic inhibitory synapses in the hippocampal CA1 region of juvenile and adult male offspring exposed in utero to anti-Caspr2 antibodies compared to controls. Additionally, local field potential (LFP) recordings from these mice show increased gamma power and altered neuronal firing patterns during social interactions, indicating functional impairments in inhibitory circuitry. These findings highlight the consequences of exposure to maternal anti-Caspr2 antibodies on PV+ interneuron development and function, providing insights into the neurobiological mechanisms underlying ASD associated behavioral phenotypes.

## Introduction

Autism Spectrum Disorder (ASD) is a complex neurodevelopmental condition characterized by deficits in social interaction and communication. Rubenstein and Merzenich proposed that a subset of ASD cases may arise from an imbalance between excitation and inhibition (E/I) in neural circuits (Lee et al., 2017; Rubenstein & Merzenich, 2003). This E/I imbalance may arise from alterations in glutamatergic and/or gamma-aminobutyric-acid (GABA) transmission, given glutamate’s role as the primary excitatory neurotransmitter and GABA as the primary inhibitory neurotransmitter, potentially contributing to the behavioral manifestations of ASD. Supporting this, postmortem brain studies from individuals with ASD reveal significant reductions in key components of the inhibitory system, such as the GABA-synthesizing enzymes (GAD67 and GAD65) and GABA receptor subunits (Blatt & Fatemi, 2011). These findings clearly implicate impaired GABAergic signaling in ASD neuropathology.

Parvalbumin-expressing (PV+) interneurons, a class of GABAergic cell originating from the medial ganglionic eminence (MGE), are characterized by their expression of the calcium binding protein parvalbumin. These interneurons are crucial for maintaining E/I balance and have been implicated in ASD and other neuropsychiatric disorders (Ferguson & Gao, 2018). Within the hippocampus, PV+ interneurons are predominantly basket cells that innervate the soma of excitatory pyramidal neurons in the CA1 region (Hu et al., 2014; Que et al., 2021). These interneurons act as pacemakers, driving network gamma oscillations that synchronize the firing of excitatory (glutamatergic) pyramidal neurons (Buzsáki & Wang, 2012). Critically, alterations in gamma oscillations, detected in frontal and temporal electroencephalogram (EEG) recordings from both children and adults with ASD, are often linked to dysfunction in inhibitory interneuron signaling (Kayarian et al., 2020). Furthermore, postmortem analyses reveal reduced PV+ interneurons and PV protein expression in the cortex of individuals with ASD compared to age-matched controls (Filice et al., 2020). A recently published RNA-Seq dataset confirmed that PV mRNA is the most strongly downregulated transcript in the cortex of subjects with ASD (Parikshak et al., 2016; Schwede et al., 2018).

Converging evidence from both environmental and genetic mouse models of ASD highlights PV+ interneuron alterations, which closely correlate with electrophysiological and behavioral deficits mirroring core ASD symptoms (Filice et al., 2020). Genetic models harboring mutations in ASD risk genes, such as contactin-associated protein-like 2 (*Cntnap2*) (Lauber et al., 2018; Thabault et al., 2024), synaptic scaffolds *Shank1* and *Shank3* (Filice et al., 2016; Gogolla et al., 2014), and phosphatase and tensin homolog (*Pten*) (Vogt et al., 2015), exhibit altered PV expression and/or PV+ interneuron numbers in key brain regions, including the hippocampus and striatum (Filice et al., 2020). Similarly, environmental ASD models, including prenatal valproic acid exposure (Gogolla et al., 2014; Lauber et al., 2016) and maternal immune activation (MIA) induced by the viral mimetic polyinosine-polycytidylic acid (Poly:IC) (Canetta et al., 2016; Vasistha et al., 2020; Yu et al., 2022), also show reductions in PV. Furthermore, PV knockout (PV^-/-^) and heterozygous (PV^+/-^) mice display core ASD-like behaviors and altered excitatory and inhibitory neurotransmission (Wohr et al., 2015). These findings emphasize that both genetic and environmental factors in ASD converge on a common mechanism involving the GABAergic inhibitory system, particularly on PV interneurons.

Caspr2, a neuronal cell-adhesion protein encoded by *Cntnap2* (ASD-risk gene) provides a compelling example of converging genetic and environmental risk factors affecting PV+ interneurons (Brimberg et al., 2016). Caspr2 expression begins during fetal development in both excitatory and inhibitory progenitor cells (Brimberg et al., 2016). Deletion of the *Cntnap2* gene (Cntnap2^-/-^) in mice results in an ASD-like phenotype, including stereotypic motor behaviors, impaired communication, and social deficits (Paterno et al., 2021; Peñagarikano et al., 2011; Peñagarikano & Geschwind, 2012). Cntnap2^-/-^ mice exhibit fewer PV+ interneurons and impaired functional connectivity (including altered gamma oscillations and sharp-wave ripple activity in hippocampal CA1) (Paterno et al., 2021). In vitro studies using interneurons cultured from Cntnap2^-/-^ mice show markedly reduced dendritic arborization (Gao et al., 2018), further highlighting the essential role of Caspr2 in the maturation and function of PV+ interneurons (Vogt et al., 2018).

Prenatal exposure to maternal anti-Caspr2 IgG also reduces PV+ interneurons in mouse offspring (Brimberg et al., 2016). Administering human monoclonal anti-Caspr2 IgG, cloned from a mother of a child with ASD, to pregnant mice at embryonic day 13.5 (E13.5), a critical time point when Caspr2 is expressed and placental antibody transfer occurs (Braniste, Al-Asmakh, Kowal, Anuar, Abbaspour, Tóth, et al., 2014), induces an ASD-like phenotype in the offspring. This includes a reduced number of parvalbumin-positive (PV+) interneurons in the CA1 region and social behavior deficits. A second, more physiologically relevant mouse model involves dams that produce endogenous polyclonal anti-Caspr2 antibodies throughout gestation (Bagnall-Moreau et al., 2020; Bagnall-Moreau et al., 2023). In this model, female mice are immunized with human Caspr2, triggering an immune response to both human and mouse Caspr2. Immunized females are then mated with non-treated male mice, resulting in dams that produce endogenous polyclonal anti-Caspr2 antibodies throughout gestation. Similar to the effects of in utero exposure to the human monoclonal anti-Caspr2 IgG, male offspring born to dams harboring polyclonal anti-Caspr2 IgG demonstrate behavioral and neuroanatomical changes, including a reduction in hippocampal PV+ interneurons and social deficits (Bagnall-Moreau et al., 2020). Interestingly, female offspring show neither structural nor behavioral alterations despite similar Caspr2 expression and exposure levels to maternal anti-Caspr2 IgG as male offspring (Bagnall-Moreau et al., 2020; Brimberg et al., 2016).

This study investigates the dysfunction of PV+ interneurons in the hippocampus following in utero exposure to maternal anti-Caspr2 antibodies. We demonstrate a selective reduction in PV+ interneurons in the CA1 region of the hippocampus of juvenile and adult male offspring exposed in utero to anti-Caspr2 antibodies, accompanied by a loss of perisomatic inhibitory synapses in the hippocampal CA1 pyramidal layer. Functionally, local field potential (LFP) recordings from the dorsal CA1 region in these mice show a significant increase in gamma power and altered neural firing patterns during a social interaction task. Together these results suggest that in utero exposure to maternal anti-Caspr2 antibodies can disrupt PV+ interneuron development, contributing to structural and functional impairments relevant to the ASD-like phenotype.

## Results

### In utero exposure to anti-Caspr2 antibody does not affect early PV interneuron development

To determine whether prenatal exposure to anti-Caspr2 IgG disrupts early PV+ interneuron development, we compared embryos and offspring from dams immunized with Caspr2 and thus exposed in utero to polyclonal anti-Caspr2 IgG (“Anti-Caspr2”) to embryos and offspring born to dams immunized with adjuvant alone and exposed in utero to nonspecific IgG (“Control”) (Bagnall-Moreau et al., 2020). Caspr2 expression begins during fetal development and is comparable between sexes throughout gestation (Brimberg et al., 2016). At E14.5, Caspr2 mRNA expression (measured by in-situ hybridization (Peñagarikano et al., 2011; Poliak et al., 1999) and transcriptional profile (Aprea et al., 2013) is localized to proliferating progenitor cells in the ventricular zone (VZ) (origin of excitatory neurons), and GABAergic progenitor cells in the medial ganglion eminence (MGE) (origin of PV+ and somatostatin interneurons) (Peñagarikano et al., 2011; Poliak et al., 1999; Vogt et al., 2018). We confirmed Caspr2 colocalization with PV+ progenitors (**Supplementary Figure 1).** Since maternal IgG enters the fetal brain between E12.5 and ∼E16.5 (Braniste, Al-Asmakh, Kowal, Anuar, Abbaspour, Toth, et al.) and Caspr2 is expressed in the MGE during this period, maternal antibodies could potentially target MGE-derived interneuron progenitors.

We examined the density of proliferating cells in the ventral MGE (vMGE), a primary source of PV+ interneurons (Hu et al., 2017; Wonders et al., 2008), using the proliferative marker phospho-histone H3 (pH3+) at E14.5 (**Figure 1A**). We found no difference in pH3+ cell density in the vMGE VZ or the dorsal MGE (dMGE) (**Supplementary Figure 2**). However, Anti-Capsr2 male embryos showed a slight increase in pH3+ cells in the SVZ compared to control males while female embryos showed no difference in the density of pH3+ cells (**Figure 1B**). Next, we assessed whether the reduction in PV (Bagnall-Moreau et al., 2020) could result from improper migration of MGE interneurons. Since PV is expressed only postnatally, we evaluated the GABA expression in cells migrating tangentially through the marginal zone (MZ) and the subventricular/intermediate zones (SVZ/IZ) from telencephalic subpallial regions where PV cells originate at E15.5 (**Figure 1C**). We found no change in the number of migrating GABA+ cells between Anti-Caspr2 and Control mice, regardless of sex (**Figure 1D**). Finally, we assessed whether exposure in utero to anti-Caspr2 IgG increased apoptosis in the MGE, contributing to the reduction of PV interneurons. TUNEL staining revealed no differences in the numbers of apoptotic cells in the MGE between Anti-Caspr2 and Control mice (**Supplementary Figure 3**). Taken together, these results indicate that early apoptosis, proliferation, and improper migration are unlikely to account for the eventual reduction in PV+ interneurons observed in Anti-Caspr2 mice.

**Figure 1:**
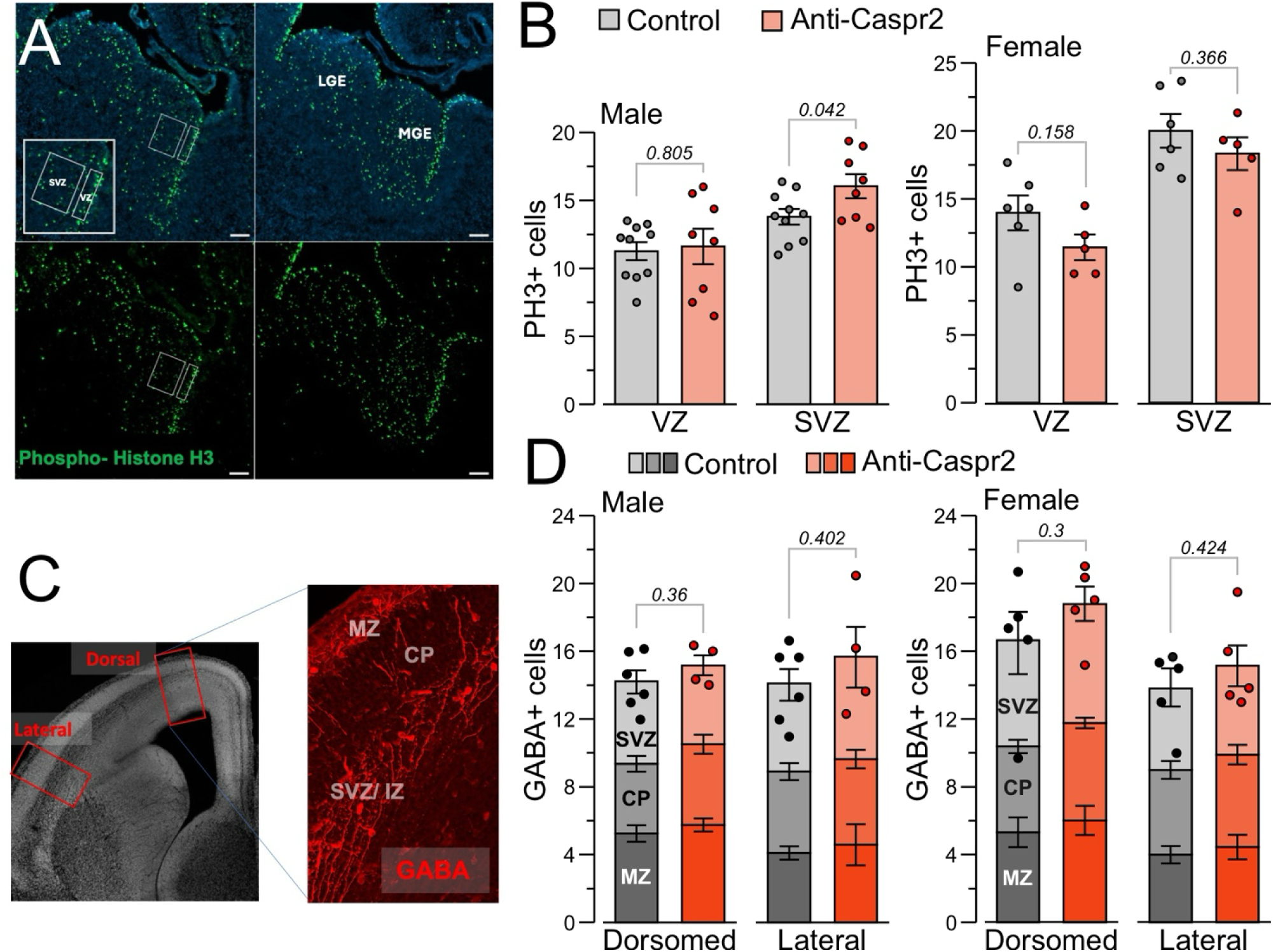
**(A)** Representative images of the phospho-histone H3 (pH3; green) staining in the MGE at E14.5. **(B)** Quantification of the number of pH3+ cells counted in the ventricular zone (VZ; 50 μm x 200 μm), and subventricular zone (SVZ; 150 μm x 200 μm) of the vMGE of Control and Anti-Caspr2 male and female embryos. Control, n = 6–10 embryos representing 3–6 litters; Anti-Caspr2, n = 5–8 embryos representing 4 litters. **(C)** Representative image of a coronal section of the embryonic telencephalon with GABA+ cells (red) at E15.5. **(D)** Stack bar plots of the number of GABA+ cells counted in 200-μm width bins in different cortical zones within the lateral or dorsal regions of the telencephalon of male and female embryos; n = 4–6 mice in each group representing 3 litters each. Data bars denote mean ± SEM; statistical comparisons with Student’s t-test. MZ, marginal zone; CP, cortical plate; IZ/SVZ, intermediate zone/ subventricular zone.

### In utero exposure to anti-Caspr2 antibody selectively reduces PV expression in male offspring

Since early developmental processes appeared unaffected, we investigated postnatal PV+ interneuron development. The reduction in PV+ interneurons could result from either decreased PV expression or the loss of PV+ interneurons, thus we examined whether the observed reduction in PV in adult mice was associated with a loss of GABAergic cells, indicative of PV+ interneuron loss. Consistent with earlier reports (Bagnall-Moreau et al., 2020; Brimberg et al., 2016), we found a significant reduction in PV+ interneuron number in the hippocampal CA1 region of Anti-Caspr2 adult males compared to control male mice. However, this significant reduction in PV+ interneurons was not accompanied by overall reduction in GABAergic cells, as the number of GABA+ cells was not different between Anti-Caspr2 and Control mice (**Figure 2B**). Notably, we observed a significant reduction in the ratio of PV+ interneurons to GABAergic cells, suggesting that rather than cell loss, PV protein expression may be downregulated at least to levels below the detection threshold of immunohistochemistry (**Figure 2A-B**). Indeed, individual PV soma fluorescence intensities were significantly lower in Anti-Caspr2 males compared to Control mice (**Figure 2C**). Finally, the reduction in PV+ interneurons was already evident in Anti-Caspr2 males at postnatal age P14 and P21 (**Supplementary Figure 4**), a defined critical period of maturation when PV is expressed in interneurons (Mukhopadhyay et al., 2009)

**Figure 2:**
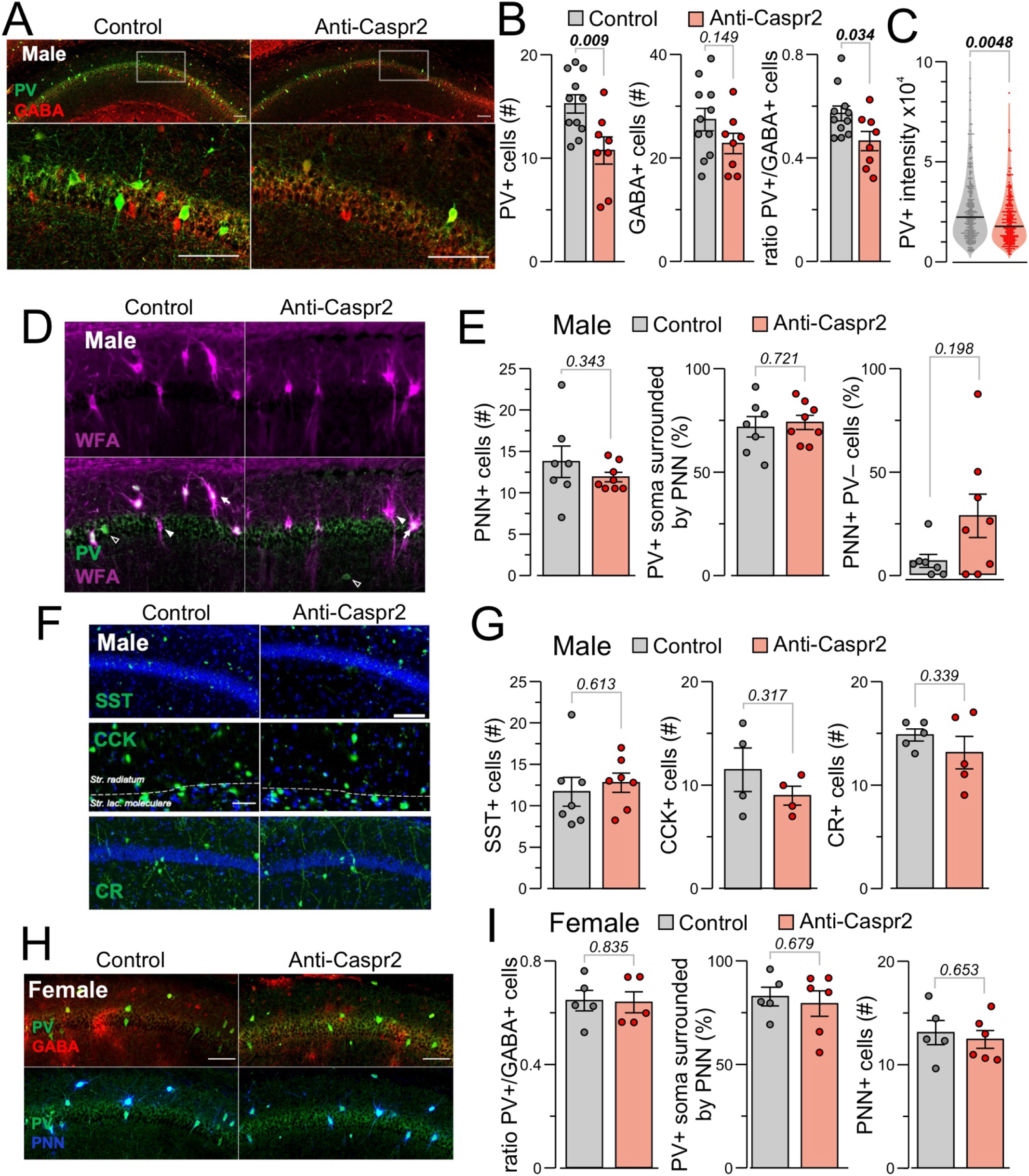
Reduction in PV expression in the hippocampal CA1 region of adult male, but not female offspring exposed in utero to Anti-Caspr2 IgG. (**A)** *Upper panel*, representative confocal images showing the expression of PV (green) and GABA (red) in the CA1 region of Control and Anti-Caspr2 male mice. *Lower panel* enlarged CA1 sections from Control and Anti-Caspr2 mice are displayed. **(B)** Quantification of the average number of cells expressing PV, GABA, and their colocalization in CA1; n = 8–11 mice per group representing 8–9 litters. (**C)** PV soma intensities are lower in CA1 of Anti-Caspr2 mice compared to Controls. Control, n = 286 cells from 6 mice representing 5 litters; Anti-Caspr2, n = 276 cells from 6 mice representing 6 litters. (**D)** Representative microscopy images of PV (green) WFA-positive PNN (magenta) in CA1 of Control and Anti-Caspr2 male mice. Arrowheads indicate PV-PNN+ cells; arrows indicate PV+PNN+ cells; empty arrow heads represent PV+PNN– cells **(E)** The average number of PNN^+^ cells in CA1 and the percentage of PV+ cells surrounded by PNNs are similar between Control and Anti-Caspr2 mice; n = 8–9 mice per group representing 8 litters. **(F)** Representative images of somatostatin (SST), cholecystokinin (CCK) and calretinin (CR) cells between Control and Anti-Caspr2 mice. (**G**) Quantification of the number of SST, CCK and CR cells between Control and Anti-Caspr2 mice. SST, n = 7 mice per group representing 4 litters; CCK, n = 4 per group representing 2 litters; CR, n = 5 per group representing 3–4 litters. **(H)** Representative confocal images showing the expression of PV+ (green) and GABA+ (red) and PNN+ (blue) cells in CA1 of Control and Anti-Caspr2 male mice. **(I)** Quantification of the number of PV+, GABA+, and PNN+ cells in CA1 between female Control and Anti-Caspr2 mice; n = 5–6 mice per group representing 5 litters. Data bars denote mean ± SEM; statistical comparisons with Student’s t-test, except (C) with Kolmogorov-Smirnov test, and (E, right) with Mann-Whitney U test.

To validate the plausible reduction in PV expression, we analyzed the distribution of perineuronal nets (PNN), specialized extracellular matrix structures that predominantly envelop the soma and proximal dendrites of PV+ interneurons (Lupori et al., 2023). PNNs can be visualized by staining with Wisteria floribunda agglutinin (WFA), a lectin that specifically binds to N-acetylgalactosamine components within the PNN structure. The presence of PNNs is strongly correlated with PV expression across multiple brain regions, specifically in CA1 (Lupori et al., 2023). We analyzed the density of PNNs in CA1 by quantifying WFA+ fluorescence intensity and the overlap with PV+ neurons (**Figure 2D**). This analysis revealed no significant differences in the average number of PNN+ cells, or the percentage of PV+ neurons surrounded by PNN (PV+PNN+) in CA1 between Anti-Caspr2 and Control males (**Figure 2E**), indicating that that the formation and stabilization of PNN onto PV+ interneurons is preserved in Anti-Caspr2 mice. The percentage of PNN+ cells surrounding cells with no PV expression (PNN+PV-cells) was trending higher in Anti-Caspr2 compared to Control mice, further supporting the hypothesis that exposure in utero to anti-Caspr2 IgG leads to reduced expression of PV rather than cell loss.

Given the lack of changes in GABA and PNN number, we considered that PV interneurons might adopt an alternative GABAergic subtype fate. Therefore, we assessed the number of other subtypes of CA1 hippocampal interneurons. We found that the reduction in CA1 PV+ interneurons was not accompanied by changes in number of other hippocampal interneurons such as somatostatin (dMGE-derived), calretinin and cholecystokinin (CGE-derived) (**Figures 2F-G**). These results suggest that in utero exposure to anti-Caspr2 IgG specifically targets PV+ interneurons, without broadly affecting other inhibitory neuronal subtypes in CA1. Furthermore, exposure in utero to anti-Caspr2 antibodies does not promote the differentiation of PV+ interneurons to an alternative GABAergic subtype fate.

Adult and juvenile female offspring did not exhibit significant differences in the number of PV+, GABA+, or PNN+ cells in the hippocampal CA1 region between Anti-Caspr2 and Control groups (**Figures 2H-I, Supplementary Figure 5**). Collectively, these findings suggest that exposure in utero to anti-Caspr2 IgG selectively impairs PV expression in male offspring, potentially altering hippocampal inhibitory circuitry and contributing to sex-specific neurodevelopmental outcomes.

### In utero exposure to Anti-Caspr2 antibody alters hippocampal CA1 network activity in male offspring

To assess the impact of in utero exposure to anti-Caspr2 IgG on neural activity and behavior, we performed LFP recordings in the CA1 region of the hippocampus and analyzed interneuron firing rates in adult male offspring. We recorded adult male Anti-Caspr2 and Control mice for three sessions (S1, S2, and S3). In S1, the recorded mouse was alone, in S2, another mouse was introduced, and recordings were conducted in a social context, and in S3, the recorded mouse was alone again (**Figure 3A**). We examined the gamma frequency band (50–90 Hz) of the LFP recordings and found that gamma power was significantly higher in Anti-Caspr2 mice across all 3 sessions (**Figure 3B, C**). We isolated the spike activity arising from putative interneurons (**Figure 3D**) and compared the mean and peak firing rates of this neural population across the 3 sessions. Remarkably, interneurons recorded from Anti-Caspr2 mice had significantly higher mean and peak firing rates during S2 (the social session), but not during S1 and S3 (the alone sessions; **Figure 3E**). To determine whether interneuron firing rates were elevated during social interactions, we analyzed firing rates during periods when the mice were in close proximity (and excluded periods when mice were distant from each other). This systematic analysis revealed significantly higher firing rates in Anti-Caspr2 mice during these periods of interaction (**Figure 3F**).

**Figure 3:**
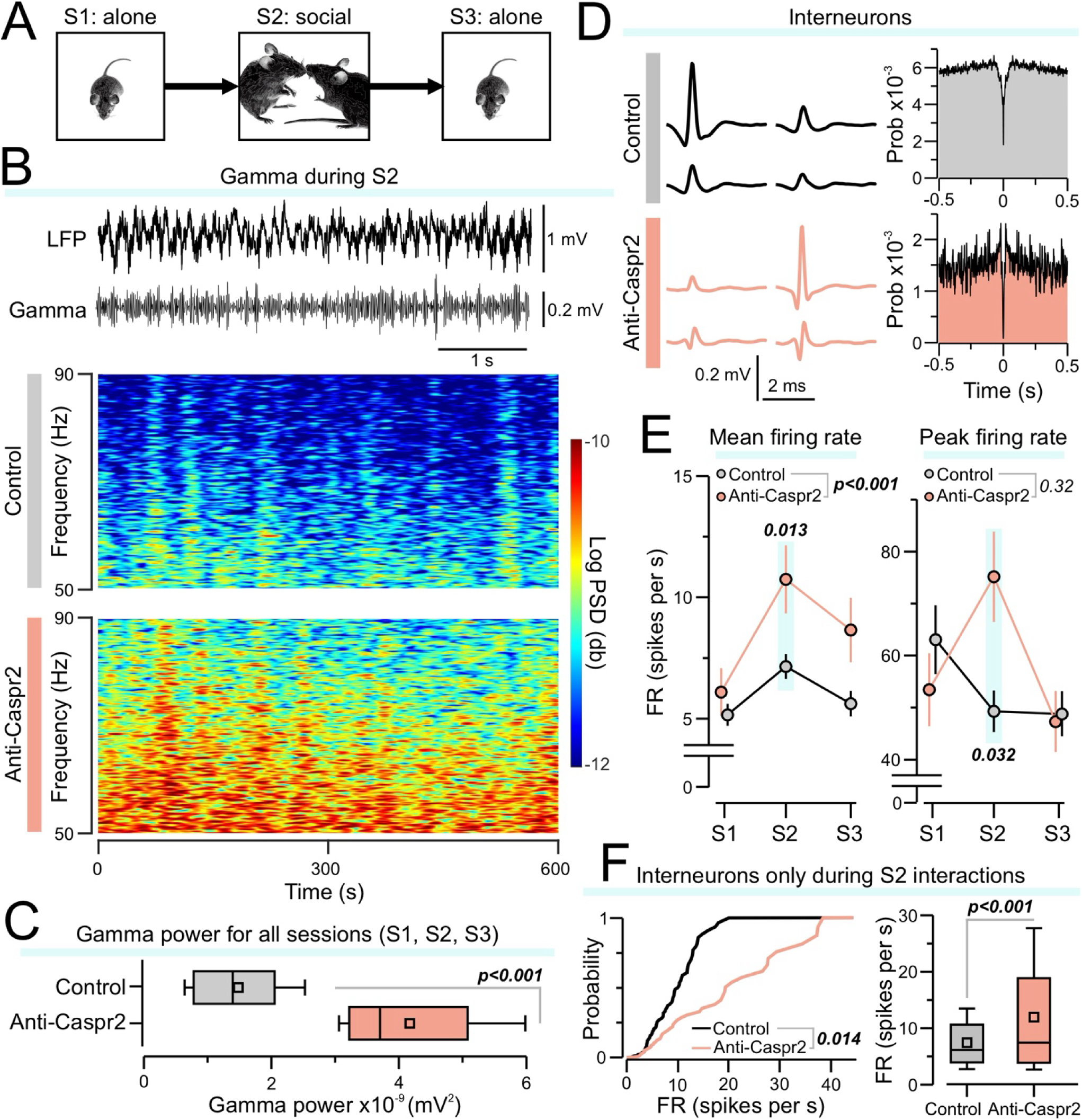
Increased power of gamma oscillations and higher interneuron firing rates in the hippocampus of the anti-Caspr2 mice. **(A)** Behavioral paradigm for recordings. Each mouse was recorded for 3 sessions (S1, S2, and S3, 10 min each), which were conducted in the same behavioral chamber. In S1, the mouse was recorded while alone. In S2, a novel animal was placed into the chamber, and they were allowed to freely interact during recordings. In S3, the mouse was recorded alone again. **(B)** *Top*, example 5-s long traces from S2 showing the LFP, and filtered gamma band. *Bottom*, example spectrograms showing the gamma power for the entire duration of S2. **(C)** Box-and-whisker plots depicting the power of gamma oscillations for S1, S2, and S3 (all sessions were combined into a single plot). The average power of the gamma band was taken for each session (P < 0.001, Kolmogorov-Smirnov test). **(D)** *Left*, average waveforms for a representative isolated interneuron, with each waveform originating from a single channel of a tetrode. *Right*, autocorrelograms for the spike times of each interneuron. **(E)** Plots showing the mean (*left*), or peak (*right*) firing rates of interneurons calculated for each recording session. Each point shows the mean ± SEM. For the mean firing rate, there were significant differences between groups across sessions (P < 0.001, two-way repeated measures ANOVA) as well as during S2 only (P = 0.013, Tukey post-hoc test). For the peak firing rate, the groups were not significantly different across sessions (P = 0.32), but there was a significant difference for S2 (P = 0.03, Tukey post-hoc test). **(F)** Cumulative probability (P = 0.014, Kolmogorov-Smirnov test) and box-and-whisker (P < 0.001, Dunn’s post-hoc test) plots depicting the firing rates for each interneuron specifically during time intervals corresponding to social interactions between mice in S2. Box-and-whisker plots show mean (square), median (line), inter-quartile range (box), and 10-90 distribution (whiskers).

### In utero exposure to anti-Caspr2 antibody reduces inhibitory synapses and PV neuropil density

Within the CA1 region of the hippocampus, PV+ interneurons are located in the stratum oriens (SO) and stratum pyramidale (SP) and primarily innervate the somata and proximal dendrites of hippocampal pyramidal cells. Therefore, we quantified inhibitory synapses in the SP of hippocampal CA1 sections that were stained for vesicular GABA transporter (vGAT), gephyrin, and NeuN. The CA1 subfield was imaged using confocal microscopy (**Figure 4A**). The fluorescent signals from synaptic proteins were converted to binary images, and the numbers of puncta were quantified in each region. We also observed reduced inhibitory synapses within the SP, as evidenced by decreased numbers of colocalized VGAT+Gephyrin+ puncta (**Figure 4B**). PV immunoreactivity in the processes of interneurons located within the SP and SR was also analyzed, revealing a significant decrease in PV+ interneuron neuropil density in Anti-Caspr2 mice (**Figure 4C–D**).

**Figure 4:**
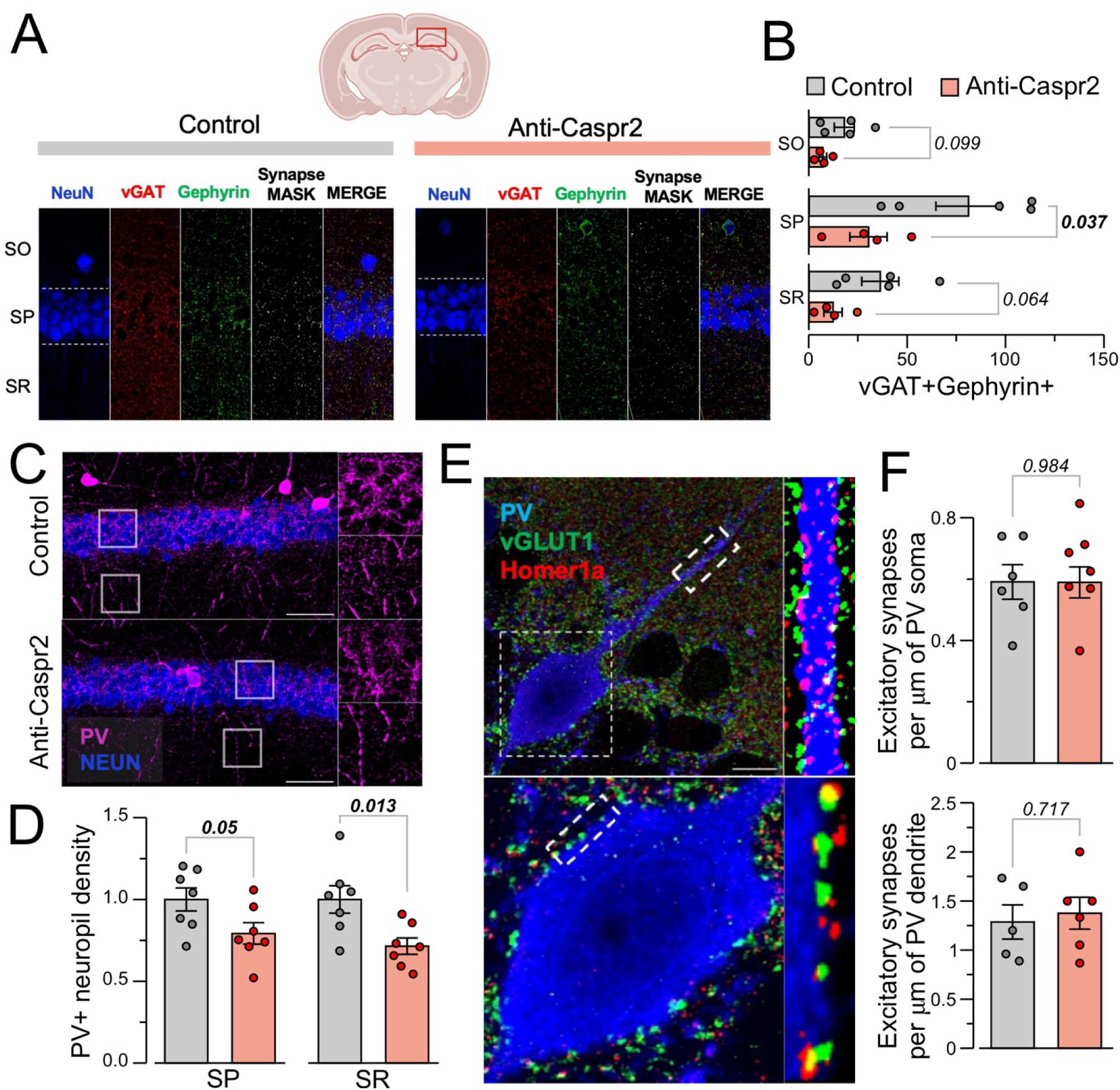
Decreased perisomatic inhibitory synapse number in the Anti-Caspr2 hippocampal CA1. **(A)** High resolution, representative images of immunofluorescent staining with NeuN (blue), presynaptic marker vGAT (red), and postsynaptic marker gephyrin (green), as well as a binary mask (white) of double positive vGAT+gephyrin+ puncta in CA1’s stratum oriens (SO), stratum pyramidale (SP), and stratum radiatum (SR). **(B)** Quantification of the density of vGAT+gephyrin+ puncta in SO, SP, and SR in Control and Anti-Caspr2 mice; n = 4–5 mice per group representing 2–3 litters; statistical comparisons with Student’s t-test with Welch correction. **(C)** Representative images showing PV+ immunoreactive neuropil in the SP and SR layers of CA1. **(D)** Relative PV neuropil density in a 50 µm x 50 µm area in the SP and SR layers. Data bars represent mean ± SEM; n = 7 mice per group representing 7 litters each, Student’s t-test. **(E)** Density of vGLUT1+Homer1a+ glutamatergic synapses onto the soma and primary dendrites of PV cells in CA1 of Control and Anti-Caspr2 adult mice. Representative images showing the distribution of vGLUT1+ (*green*) and Homer1a+ (*red*) synaptic puncta on the soma (bottom panel) and dendrite of a PV^+^ neuron. **(F)** Quantification of the number of excitatory synapses (vGLUT1+Homer1a+ colocalized puncta) around the soma and primary dendrites (15-µm length) of PV+ neurons; n = 5–6 mice per group representing 4 litters. Data bars denote mean ± SEM; Student’s t-test.

CA1 interneurons receive excitatory glutamatergic input from neighboring pyramidal neurons, as well as from extrahippocampal cortical and thalamic afferents (Booker & Vida, 2018), which are critical for proper development and maintenance. A reduction in PV expression and the altered function of these interneurons could reflect changes in excitatory input onto the inhibitory cells (Patz et al., 2004). Using Airyscan super-resolution microscopy, we imaged synapses on the soma and the proximal dendrites of PV-expressing cells located in the CA1 SP subregion immunolabelled for the detection of the synaptic proteins vGLUT1 (pre-synapse) and Homer1 (post-synapse) (**Figure 4E**). Overall, there were no significant changes in the density of overlapping vGLUT1+Homer1+ puncta on the soma or proximal dendrites of PV+ interneurons (**Figure 4F**).

Altogether our results demonstrate that hippocampal CA1 PV interneurons in Anti-Caspr2 mice exhibit reduced PV expression, decreased inhibitory synapses, and reduced neuropil density, despite no significant changes in the density of excitatory synaptic inputs onto their soma or proximal dendrites. These findings suggest that mechanisms beyond glutamatergic input, including intrinsic cell changes or broad network-level disruptions, may contribute to the observed deficits in PV expression and function.

## Discussion

Both genetic and environmental perturbation to Caspr2 during in utero development can lead to neurodevelopmental disorders, including ASD (Brimberg et al., 2016; Rodenas-Cuadrado et al., 2014; Rodenas-Cuadrado et al., 2016; St George-Hyslop et al., 2022). This study demonstrates that exposure in utero to anti-Caspr2 IgG specifically disrupts PV expression in the hippocampal CA1 region of male offspring, without affecting the overall number of GABAergic interneurons. This selective reduction in PV expression leads to impaired inhibitory signaling and altered network dynamics, potentially contributing to the behavioral phenotypes characteristic of ASD.

Exposure in utero to anti-Caspr2 IgG led to selective loss of PV expression within the hippocampal CA1 region of male mice offspring. The preferential vulnerability of PV interneurons compared to other GABAergic interneurons in Anti-Caspr2 mice may be influenced by several factors, such as the timing of maternal antibody transfer across the placenta (and into the developing brain), the level of Caspr2 protein expression during that critical window, or a combination of both factors. Somatostatin interneurons are among the earliest born MGE-derived progenitors, with peak generation occurring earlier than the PV interneurons which are generated over a broader window extending into late embryonic stages (Butt et al., 2005). The overlap with the timing of antibody exposure may contribute to the specific impact of anti-Caspr2 IgG on MGE-derived PV+ interneurons. Additionally, *Cntnap2* transcript levels are significantly lower in somatostatin interneurons compared to PV+ interneurons in P28–P35 mouse brains (Vogt et al., 2018), further supporting the selective effect on PV interneurons. (Bocchio et al., 2020; Pelkey et al., 2017).

We observed reduced PV expression as early as P14 and P21, a critical developmental window for PV+ interneuron maturation (Hu et al., 2014). Importantly, this reduction in PV does not appear to arise from disrupted key developmental processes, including the proliferation or migration of MGE-derived progenitors, apoptotic cell death, or shifts in interneuron subtype specification, as indicated by observations during embryonic and early postnatal development. One limitation of our data is the lack of molecular markers specific to PV in the developing brain, which presents a significant challenge in identifying putative PV interneurons prior to the onset of expression. While studies have suggested markers to stratify PV interneurons at the embryonic (Allaway et al., 2021; Bandler et al., 2017; Pai et al., 2019) and early postnatal age (Pai et al., 2019), these markers lack a distinct and specific expression pattern exclusive to PV.

The detection of PV in our study, as in others, primarily relies on immunostaining with anti-PV antibodies. However, using both GABA and PNN as surrogate markers we confirmed the presence of cells that have likely lost parvalbumin immunoreactivity. Postmortem brain studies of individuals with ASD have revealed a reduction in PV+ interneurons (Ariza et al., 2018; Hashemi et al., 2017). Multiple ASD mouse models, including the Caspr2 deficient mouse, recapitulate the reduced PV+ interneuron density in various brain regions including the hippocampus (Filice et al., 2016; Lauber et al., 2018). However, those studies did not determine whether the observed reductions reflect an actual loss of cells or a decrease in PV protein expression falling below the detection threshold (Lauber et al., 2016, 2018). This distinction has important implications as it likely will suggest different therapeutic modalities.

PV expression levels have been shown to be dynamic in response to learning and sensory experience (Miranda et al., 2022; Patz et al., 2004). Indeed, it is possible, that reduced PV expression is the result of reduced excitatory synaptic input or activity as this has been observed in patients with schizophrenia (Chung et al., 2016). Consistent with this, structural changes have been found on the dendrites of CA1 excitatory pyramidal neurons in Anti-Caspr2 mice (Bagnall-Moreau et al., 2020), and a reduction in synaptic connectivity and density is observed in Caspr2 deficient mouse mice (Lazaro et al., 2019). However, we have not found differences between adult Anti-Caspr2 and Control male mice in the density of excitatory synaptic input on PV+ interneurons in the pyramidal layer of CA1. Taken together, our data suggest that the observed reduction in PV+ interneurons in Anti-Caspr2 mice is likely attributable to a loss of PV expression. It is plausible that other non-cell-autonomous factors influence PV+ interneuron maturation postnatally, underscoring the need for further investigation into these potential mechanisms. This study further highlights the importance of distinguishing between changes in protein expression and actual cell loss, as this distinction has significant implications for understanding the underlying mechanisms and developing targeted therapeutic strategies for ASD.

We found that exposure in utero to anti-Caspr2 antibodies led to hyperexcitability of PV interneurons during social test. This is in line with the study by Deng et al. 2019 showing that hyperexcitation of CA1-PV interneurons impaired social discrimination, leading to familiar mice being perceived as novel (Deng et al., 2019). The same study also found that stimulating CA1-PV interneurons at a gamma frequency impaired social recognition and created a preference for the stimulation-paired mouse. Our data also align with this study showing overall increase in gamma oscillations in mice exposed in utero to anti-Caspr2 IgG. Previous studies of freely moving Caspr2-deficient mice during spatial discrimination and social behavior tasks demonstrate alterations in CA1 and prefrontal theta and gamma rhythmicity (Mohapatra et al., 2024; Paterno et al., 2021). Gamma oscillations play a crucial role in sensory processing and cognition, facilitating the coordination of neural activity across brain regions. Alterations in gamma band activity have been consistently observed in various neurodevelopmental disorders, suggesting a common mechanism underlying cognitive deficits. Our results not only emphasize the critical role of PV interneurons in coordinating gamma oscillations but also suggest that hippocampal PV interneuron dysfunction may contribute to the behavioral and cognitive deficits observed in ASD.

Critically, we observed a significant reduction in vGAT+ synapses in the CA1 pyramidal layer of anti-Caspr2 mice. This decrease in perisomatic-targeting PV basket cell synapses suggests a disruption of inhibitory circuitry. While we found no differences in the excitatory drive onto PV+ interneurons, future studies should directly measure PV+ interneuron function and assess the intrinsic excitability of these cells in Anti-Caspr2 mice. Additionally, investigating the inhibitory input onto PV+ interneurons by measuring synaptic proteins such as vGAT, bassoon, and gephyrin would provide valuable insights into the mechanisms underlying the observed alterations in gamma oscillations and neuronal synchronization. By further elucidating the impact of exposure in utero to maternal anti-Caspr2 antibodies on PV+ interneuron function and network dynamics, we can better understand the neurodevelopmental consequences of in utero exposure to these antibodies and their potential contribution to the pathogenesis of disorders like ASD.

Consistent with the reported male bias in ASD, our model demonstrated that male offspring exposed in utero to anti-Caspr2 antibodies exhibited ASD-associated behaviors and neuroanatomical changes, whereas female offspring did not (Bagnall-Moreau et al., 2020; Brimberg et al., 2016). While ASD is more prevalent in males than females, the neurobiological mechanisms underlying these sex differences remain poorly understood. In the present study, we have focused on PV+ interneurons, as there is growing evidence for GABAergic system dysfunction in ASD. Sex differences in GABA circuit development, particularly the timing of the postnatal GABA shift, are well-documented (McCarthy et al., 2002; Peerboom & Wierenga, 2021). This transition from depolarizing to hyperpolarizing effects occurs later in males than in females in the hippocampus and is delayed in animal models of several neurodevelopmental disorders (Fernandez et al., 2019; Lozovaya et al., 2019; Roux et al., 2018). Investigating whether the developmental GABA shift impacts sex-specific PV+ interneuron maturation and connectivity could uncover critical mechanisms underlying ASD. Alternatively, gonadal hormones play a modulatory role in neuronal development, contributing to sex-biased behaviors and traits (Bordt et al., 2024). Estrogen receptor β, for instance, is highly colocalized within PV+ interneurons in the hippocampus, suggesting that hormones such as estradiol may mediate organizational effects on neuronal maturation (Blurton-Jones & Tuszynski, 2002; Wu et al., 2014). Further studies are needed to explore whether sex-specific differences in other GABA signaling components during brain development contribute to distinct cognitive and behavioral outcomes (Ethiraj et al., 2021; Pandya et al., 2019). These differences may underlie susceptibility to risk factors, including maternal antibodies, that influence ASD pathogenesis.

In conclusion, our findings demonstrate that in utero exposure to maternal antibodies can induce structural and functional changes in PV+ interneurons within the hippocampal CA1 region of male offspring. Future research will focus on uncovering the mechanisms regulating PV expression and maturation to further understand their role in interneuron function and social behavior. Additionally, investigating how sex differences influence developmental changes in PV interneurons and the impact on hippocampal network dynamics could provide critical insights for developing targeted therapies aimed at addressing specific cognitive domains associated with ASD symptoms.

## Materials and Methods

### Animals

All procedures conformed to the National Institutes of Health Guidelines for the Care and Use of Laboratory Mice and were approved by the Feinstein Institutes Animal Care and Use Committee (IACUC). Female and male C57BL/6 mice (5 weeks old) were obtained from the Jackson Laboratory. After one week of acclimation, female mice were randomly assigned to be immunized in either adjuvant alone or with Caspr2.

### Immunization and timed pregnancies

As described previously (Bagnall-Moreau et al., 2020; Rubio-Marrero et al., 2016), female mice were immunized intraperitoneally (i.p.) with 50 µg (in 50 µl of saline) of the extracellular region of human Caspr2 expressed using glycosylation deficient HEK293T GnTI − cells (Rubio-Marrero et al., 2016), in Complete Freund’s Adjuvant (50 µl). Subsequent booster injections of immunogen (50 µg in 50 µl of saline) mixed with Incomplete Freund’s Adjuvant (50 µl) were administered on day 14 and 28. Titers to both human and mouse Caspr2 were determined by live cell based assay as described in (Bagnall-Moreau et al., 2020). Two weeks following the last boost, immunized female mice were mated with naïve, untreated male mice. To generate timed pregnancies, a pair of females were housed with a single male at the start of the dark cycle for 14 hours. We designated the first day as Embryonic day (E) 0.5. Whole embryos or dissected brains were harvested from E14.5 to E18.5, and processed for sex identification, blood collection, and fetal brain pathology. Additional pregnancies were carried to full term, and brains were harvested after perfusion-fixation from postnatal and adult offspring for histological assessments.

### Tissue preparation and immunohistochemistry

Whole embryos (E14.5) or dissected brains (E15.5) were harvested, briefly rinsed in ice-cold HBSS, and immersion-fixed in 4% paraformaldehyde (PFA) prepared in 0.1 M phosphate buffer (PB) for 24 hours at 4°C. After fixation, the tissues were transferred through graded sucrose concentrations for cryoprotection, embedded in optimal cutting temperature (OCT) compound, and frozen over dry ice. Cryosections were prepared at 14 µm using a Leica CM1850 cryostat. Alternatively, to detect GABA immunoreactivity, embryonic brains were removed, and immersion fixed in 4% PFA / 0.2% Glutaraldehyde in 0.1 M PB solution for 24 hr. at 4°C. After several washes in phosphate buffered saline (PBS), fixed brains were cut at 50 microns in ice-cold PBS on a vibratome (Leica VT 1200S). Postnatal and adult offspring were deeply anesthetized with Euthasol before transcardial perfusion with heparinized saline followed by 4% PFA in

0.1 M PB. Brains were dissected and postfixed overnight in 4% PFA, then cryoprotected in 30% sucrose in 0.1M PB at 4°C. Free floating sections (30 μm) were prepared on a sliding microtome and placed in ethylene glycol storage solution at 4°C until use. Alternatively, for the detection of GABA, postnatal and adult mice were perfused with a fixative solution containing 4% paraformaldehyde (PFA) and 1% glutaraldehyde in 0.1M PB followed by post-fixation in the same solution for 1 hour. After several washes in PBS, brains were cut at 30 µm thickness in ice-cold PBS on a vibratome, followed by reduction in 1% sodium borohydride in 0.1M PB at room temperature for 30 min and several PBS washes. Sections were stored in PBS with 0.05% sodium azide until use. To account for differences in hippocampal region sizes between groups of adult mice, sections at the coronal level that corresponds to bregma AP −1.70 to −2.06 was selected for dorsal hippocampus and AP −2.70 to −2.80 for ventral hippocampus (Paxinos & Franklin, 2019). For free floating immunofluorescence, non-specific binding in sections was blocked with 3% normal goat serum (Gibco) with 0.3% Triton X-100 in PBS for 1h at RT. Primary antibodies were added and incubated overnight at 4 °C with gentle rocking. Sections were then washed in PBS with 0.2% Tween-20 (PBST) and incubated at 1.5 hr. at room temperature with the appropriate fluorophore conjugated secondary antibody. Biotinylated wisteria floribunda agglutinin (WFA) was used for the detection of perineuronal nets and was visualized using fluorophore-conjugated streptavidin. After the sections were washed with PBS, they were stained with the nuclear marker 4,6-diamidino-2-phenylindole (DAPI, 1µg/ml) for 5 min. After a final wash in PBS, sections were mounted onto glass slides with coverslips using Fluorescent Mounting Medium (Dako). Negative controls were performed for each primary antibody.

Immunofluorescence experiments were performed using the following primary antibodies: anti-parvalbumin (mouse, MAB1572, Millipore), anti-parvalbumin (rabbit, 195-002, Synaptic Systems) anti-gephyrin (mouse, 147011, Synaptic Systems) anti-cleaved caspase 3 (rabbit, 9661S, Cell signaling technologies), anti-vGLUT1 (guinea pig, AB5905, Millipore), anti-vGAT (guinea pig, 131004, Synaptic Systems), anti-CCK8 (rabbit,C2581-25UL,Millipore), anti-homer (rabbit, 160-003, Synaptic Systems), anti-LHX6 (mouse, sc-271433, Santa-Cruz), anti-GABA (rabbit, AB5016, Sigma), anti-somatostatin (rat, MAB354, Fisher Scientific), anti-calretinin (rabbit, 214 102, Synaptic Systems), Biotinylated Wisteria Floribunda Agglutinin (NC1042310, Fisher Scientific), anti-phospho histone H3 (rabbit, 06-570, Millipore).

### RNAscope in situ hybridization

RNA fluorescence in situ hybridization (RNA FISH) was performed in a single experiment, or in combination with immunohistochemistry on 14 µm cryosections of whole embryos, or coronal brain slices using the RNAscope Multiplex Fluorescent Reagent Kit v2 assay (Advanced Cell Diagnostics) according to the manufacturer’s instructions. Hybridization was performed with RNAscope probes Mm-LHX6, Mm-NKX21, Mm-CNTNAP2. Nuclei were stained with DAPI (1µg/ml). Images were acquired under a Zeiss LSM 880 confocal microscope.

### TUNEL Assay

For the detection of apoptotic cells at E14.5, sections were prepared and cut as described above. A TdT-mediated dUTP nick-end labeling (TUNEL) assay was performed on embryonic cryosections using the Fluorescein In Situ Cell Death Detection Kit according to the manufacturer’s protocol (Roche).

### Image Acquisition and Analysis

All fluorescent and brightfield images were obtained with a Zeiss apotome 2 or on a Zeiss LSM 880 confocal Axio observer microscope. For postnatal and adult histology (PV, SST, CR, CCK, WFA, GABA, NEUN), 4-8 z-stack images from 3-4 free floating sections of both hemispheres of the hippocampus were acquired using a Plan-Apochromat 20x*/* NA 0.8. For super-resolution imaging of synaptic proteins (vGAT, vGLUT1, Gephyrin, Homer1a), images were collected using a Zeiss LSM 880 microscope (ZEN 2.3 black edition, software v.14.0.9.201) equipped with plan-apochromat 63x oil-immersion objective (1.4 NA) and Airyscan Gallium Arsenide Phosphide (GaAsP) detector. ZEN ‘.czi ’ files were processed into 8-bit or 16-bit TIFF formats for analysis in ImageJ.

### Cell Quantification

The Cell Counter ImageJ plug in was used to quantify the number of cells within each image. For the analysis of proliferating cells in MGE E14.5 samples, the number of pH3+ cells in the VZ surface (50 µm x 200 µm) and SVZ (150 µm x 200 µm) of the dorsal MGE in sections at equivalent rostro-caudal locations was measured. To analyze GABAergic cell distribution at E15.5, GABA-immunoreactive cells, identified by soma staining, were quantified within 200 µm-wide bins across lateral and dorsomedial regions of the embryonic cortex. For cell counting of interneuron markers in the hippocampal CA1 region of postnatal and adult mice, acquired images were first converted to maximum projection. To analyze interneurons in postnatal and adult mice, z-stack images were converted into maximum projections, and the number of cells expressing the interneuron marker was quantified in the CA1 region of the dorsal hippocampus.

### Synaptic protein quantification

For the analysis of GABAergic synapses in the hippocampal CA1 region, 30 µm coronal brain sections containing the dorsal hippocampus were stained with presynaptic (vGAT) and postsynaptic (gephyrin) markers. Tiled z-stack images with a 0.25 µm step size were acquired and analyzed using ImageJ. Local background subtraction was performed using the rolling ball method, and binary masks were generated for each channel with optimized settings applied uniformly across all images. The ImageJ ‘AND’ operation was applied to identify colocalized puncta between the masks of presynaptic and postsynaptic signals, enabling the quantification of inhibitory synapses The analysis of synaptic contacts was performed using a modified macro based on the method described (Guirado et al., 2018). Briefly, multi-channel images were subjected to post-processing methods including background subtraction and Gaussian blur filtering. Regions of interest (ROIs) were defined around PV cells to measure the perimeter of the soma or dendrite, which was then enlarged by 1 µm to create a ‘mask’ for analyzing synaptic signals within the perisomatic region. Intensity thresholds were determined and binarized for each synaptic marker channel. The ‘Watershed’ and ‘Analyze Particle’ tools were subsequently used to quantify presynaptic puncta (vGLUT1, size ≥ 0.1μm²), postsynaptic puncta (homer1a, size ≥ 0.1 μm²), and colocalized vGLUT1+/homer1a+ synaptic contacts on PV somas or dendrites.

### Neural recordings in the hippocampal CA1 region of freely behaving mice

Custom-designed microdrive bodies were 3D-printed (Form-3, Formlabs, Somerville, MA). Each microdrive was designed to carry four tetrodes, for a total of 16 channels. We attached polyimide tubing and an electrode interface board (Omnetics EIB-16, Neuralynx, Bozeman, MT) to each microdrive body. Tetrodes were wound from 90% platinum, 10% iridium wire (17.8 µm diameter; California Fine Wire, Grover Beach, CA) and threaded through the polyimide tubes to create a movable array (Chang et al., 2013). On the day of implantation surgery, we cut the tetrodes at a ∼45° angle. This ensured that, when implanted, the lowest tetrode was positioned in stratum pyramidale and the other tetrodes were located in stratum oriens. We then electroplated the tetrodes with platinum black solution (Neuralynx) to achieve an impedance under 300 kΩ for each tetrode. All surgical procedures were performed under isoflurane anesthesia. Before incision, mice received buprenorphine (0.1 mg/kg) and meloxicam (5 mg/kg) for pain relief. We shaved the fur from the surgical site, which was then scrubbed with betadine and isopropyl alcohol. After making an incision and exposing the skull, we applied a layer of C&B-Metabond (Parkell, Edgewood, NY) and allowed it to dry. Two craniotomies were made, one over the cerebellum and the other over the hippocampus. The coordinates used for dorsal CA1 were [AP, −2.18, ML, −1.5] from bregma. We installed the ground screw into the craniotomy located in the occipital bone. Then, the microdrive was aligned so that the tetrodes were directly above the intended region and secured with dental acrylic. Implanted mice were observed for three days post-surgery and received daily meloxicam injections (5 mg/kg). Over the next three days, tetrodes were lowered until the lowest one reached stratum pyramidale, identified by the appearance of theta waves and bursting spike activity characteristic of CA1 pyramidal neurons.

Neural recording during behavioral tasks was performed 7–10 days post-surgery. Recordings consisted of three 10-minute sessions: alone (S1), with another mouse present (S2), and alone again (S3). After acquisition, data were transferred to a Linux machine (Ubuntu 18.04.5 LTS) for automated spike sorting with MountainSort (Strohl et al., 2021; Chung et al., 2017) and analyzed with NeuroExplorer version 5 (Plexon, Dallas, TX). LFP data were analyzed with custom-written MATLAB scripts and the Chronux toolbox (http://chronux.org/; Mitra, P., & Bokil, H. (2008). Observed Brain Dynamics. Oxford University Press) on a Windows 10 PC. Final results were processed in MATLAB, Excel (Microsoft), and Origin 2024 (OriginLab, Northampton, MA).

To confirm the location and positioning of the electrode, we performed Cresyl violet counterstaining on implanted mice as described in [(ref).

### Statistics

Statistical analysis of the data was performed with Origin. Normally distributed data were analyzed using the student’s t-test (for two groups) or by one-way ANOVA test (for more than two groups). We used Mann– Whitney test to compare medians of two independent groups for small datasets or for non-normal distributed samples.

## Acknowledgements

We would like to thank Dr. Betty Diamond for helpful discussions. The study is supported by a NIH grant R21MH136489A (given to LB) and a postdoctoral fellowship from Autism Speaks 12996 (given to CBM). LB and CBM are also the recipient of Advancing Women in Science and Medicine (AWSM) awards. BS is supported by NIH T32 5T32AI155392.

## Author contributions

CBM, JS, PH, and LB designed the experiments. CBM, JS, CC, BS, PH, and LB. performed the experiments. CBM, JS, PH, and LB analyzed the data. CBM, PH and LB wrote the paper.

## Supplemental Information

**Supplemental Figure 1:**
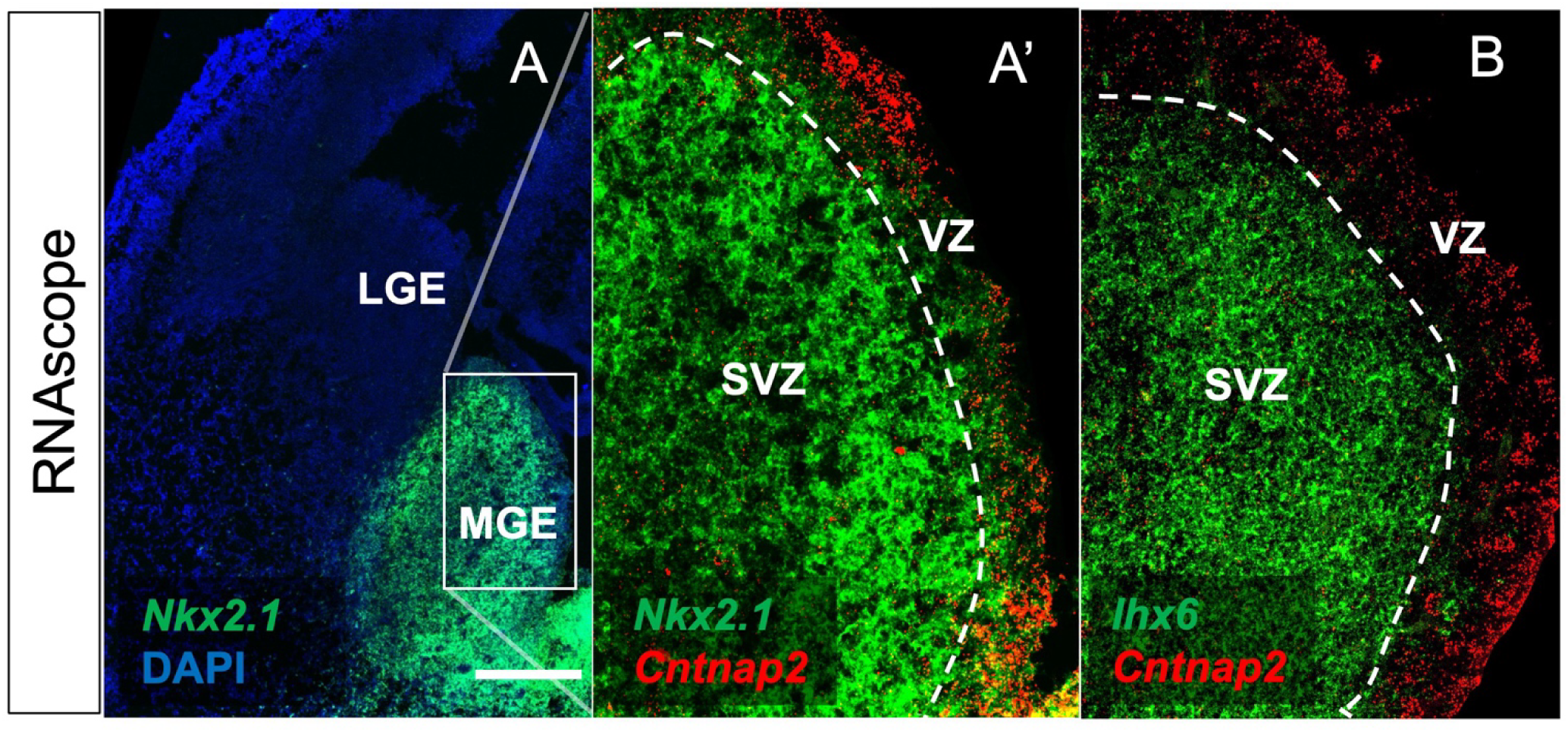
Cntnap2 is expressed in the VZ of the MGE at E14.5 (C57BL/6 wild type). RNAscope dual fluorescence in situ hybridization of Cntnap2 (in red) is colocalized with Nkx2.1 (in green, panel **A’**), but not with Lhx6 (in green, panel **B**) in the MGE. MGE, medial ganglionic eminence; VZ, ventricular zone; SVZ subventricular zone. Scale bar in **A**: 250 µm.

**Supplemental Figure 2:**
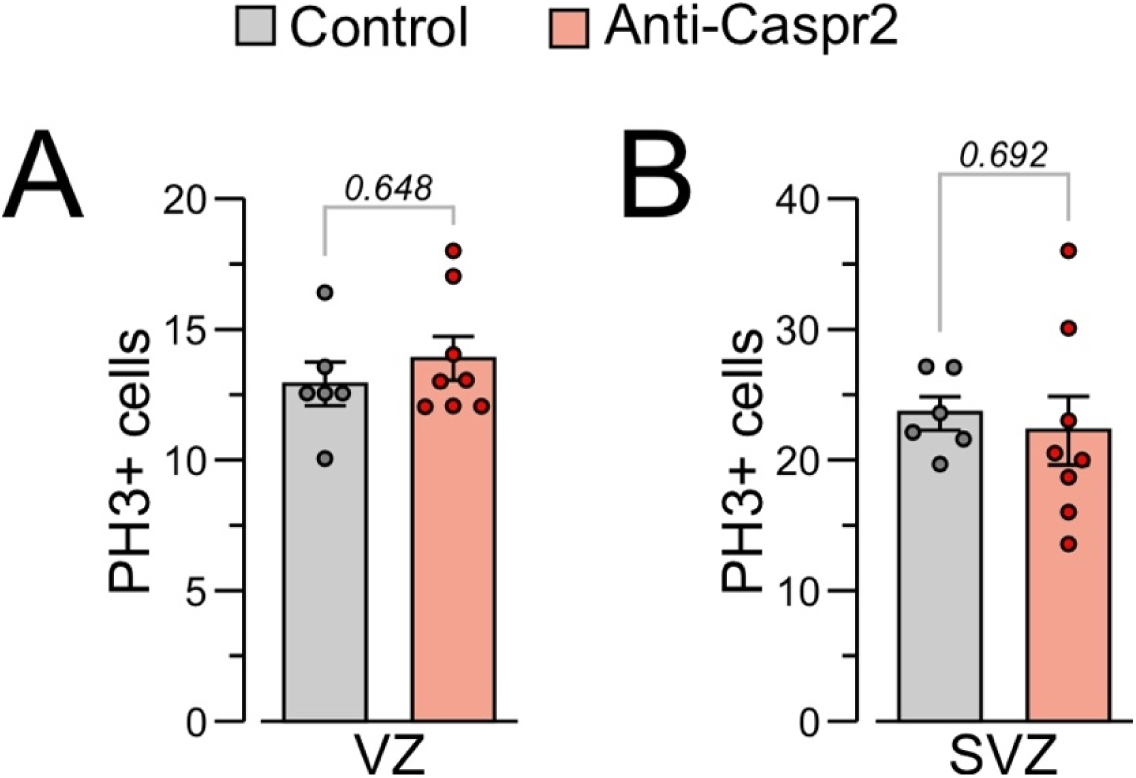
(**A**) Quantification of the number of pH3+ cells counted in the ventricular zone (VZ; 50 μm x 200 μm) and (**B**) subventricular zone (SVZ; 150 μm x 200 μm) of the dMGE of Control and Anti-Caspr2 E14.5 male embryos. Control, n = 6 embryos representing 3 litters; Anti-Caspr2, n = 8 embryos representing 4 litters. Data bars denote mean ± SEM; statistical comparisons with (A) Mann-Whitney U test and (B) Student’s t-test.

**Supplemental Figure 3:**
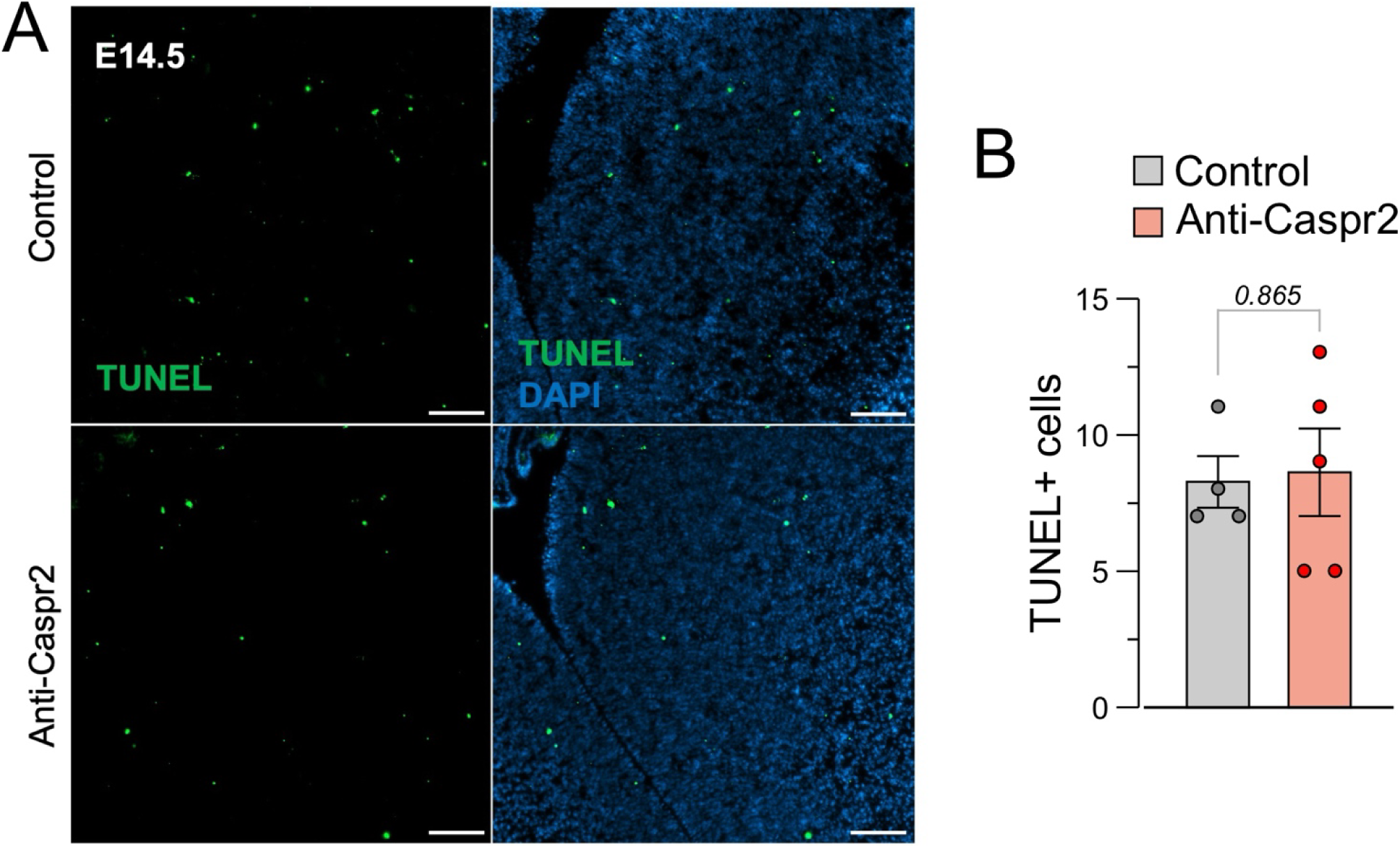
No differences in the number of apoptotic cells in the MGE. (**A**) Representative images of the TUNEL+ staining in MGE (green) at E14.5. Scale bar = 50 μm. (**B)** Quantification of the number of TUNEL+ cells in the MGE of Control and Anti-Caspr2 E14.5 male embryos. n= 4–5 mice representing 2–3 litters. Data bars denote mean ± SEM; Student’s t-test.

**Supplemental Figure 4:**
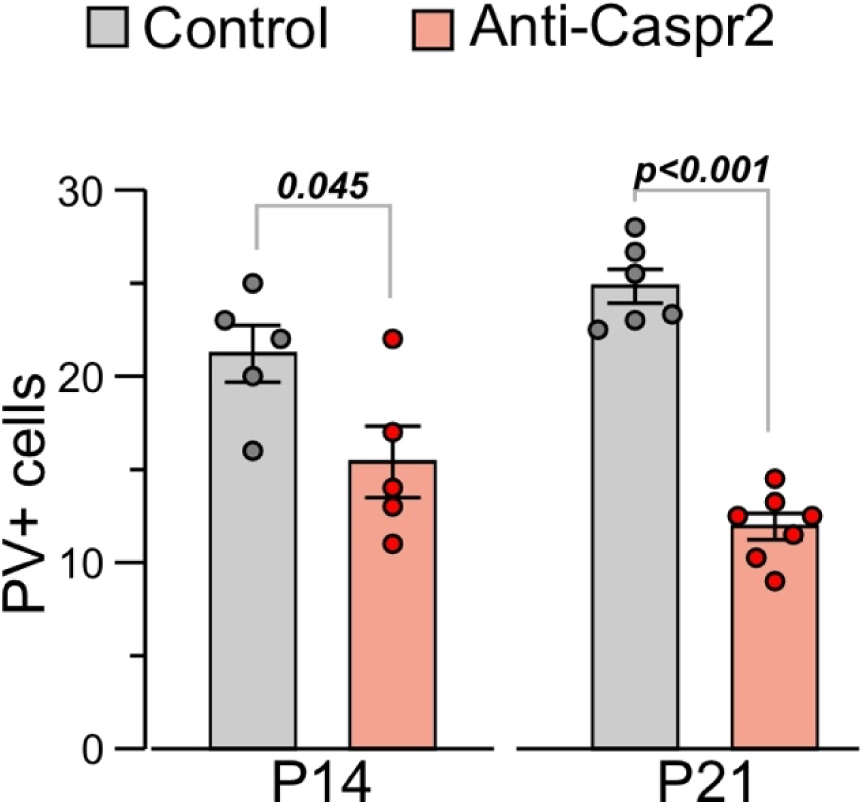
Reduction in PV expression in P14 and P21 juvenile male mice. Number of PV+ interneurons counted in CA1 of Control and Anti-Caspr2 mice on postnatal days P14 and P21; n = 5–6 mice per group representing 3 litters each. Data bars denote mean ± SEM; Student’s t-test.

**Supplemental Figure 5:**
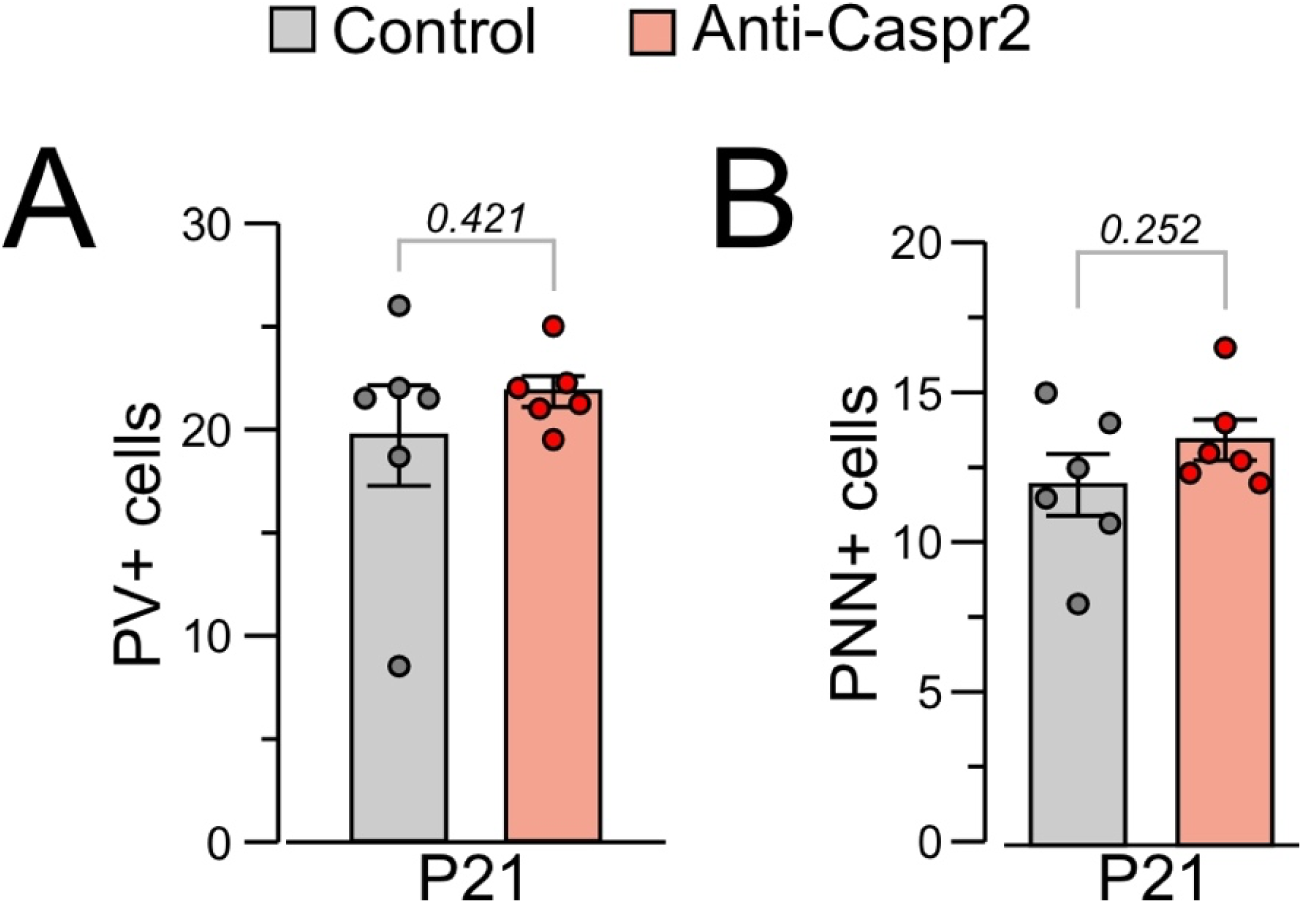
No differences in the number of PV+ or PNN+ cells at postnatal day 21 between female Control and Anti-Caspr2 mice; n = 6 mice per group representing 4–5 litters. Data bars denote mean ± SEM; Student’s t-test.

